# rTCT: Rodent Triangle Completion Task to facilitate translational study of path integration

**DOI:** 10.1101/2025.02.04.636463

**Authors:** Stephen Duncan, Sulaiman Rehman, Vladislava Sagen, Irene Choi, Sami Lawrence, Om Kalani, Lisette Gold, Lillian Goldman, Sophia Ramlo, Kylene Stickel, Dylan Layfield, Thomas Wolbers, Zoran Tiganj, Ehren L. Newman

**Affiliations:** Department of Psychology & Brain Sciences, Indiana University; 1101 E. 10^th^ Street, Bloomington, IN, 47405, U.S.A.; German Center for Neurodegenerative Diseases (DZNE), 39120, Magdeburg, Germany

## Abstract

Path integration is navigation in the absence of environmental landmarks and is a primary cognitive mechanism underlying spatial memory. Path integration performance is primarily assessed in humans using the Triangle Completion Task (TCT). In humans, TCT has shown promise for the early diagnosis of Alzheimer’s disease. In rodents, however, path integration is assessed using a wide variety of tasks but none of which currently provide a homologue for the TCT. As rodents are routinely used as preclinical models, homologous path integration tasks that result in comparable performance metrics between species are important. In the present study we developed and tested a novel rodent version of the triangle completion task to enhance cross species comparability of path integration performance. Rats were able to comprehend and perform the task. A group of Alzheimer’s disease model rats (TgF344-AD) exhibited similar path integration performance to their wild-type littermates; however, analysis of behavioural structure suggests use of differing behavioural strategies. This work establishes a novel rodent homologue of the triangle completion task, facilitating enhanced reverse translational study of human path integration.

## Introduction

Spatial memory enables animals to effectively navigate through their environment, playing a crucial role in their survival. One of the primary cognitive mechanisms underlying spatial memory is path integration, involving the use of self-motion cues to navigate to goal locations in the absence of environmental landmarks (Etienne & Jeffery, 2004). In humans path integration is most commonly assessed using the Triangle Completion Task (TCT; Segen et al., 2022) and has already shown promise for detecting Alzheimer’s disease (AD) in its earlier stages (Howett et al., 2019; Newton et al., 2024). The regular use of TCT for assessing path integration in humans, and its potential for clinical application, make the TCT a valuable behavioural assessment tool in pre-clinical research. There is, however, no homologous task in preclinical models of AD - i.e. rodents. In the present study we developed and tested a novel rodent version of the TCT.

The TCT is a widely used test of path integration in humans (Allen et al., 2004; Harris & Wolbers, 2012; Loomis et al., 1993; Mahmood et al., 2009; Marlinsky, 1999; Nico et al., 2002; Stangl et al., 2020). During this task, participants are guided from location A to location B and then to location C before being asked to take the shortest path back to A, without assistance from environmental markers. Participants must, therefore, use the idiothetic information gathered during the guided portions of the task, to navigate back to the start location.

The utility of the TCT for assessing path integration performance to advance the mechanistic understanding of mental health has been clearly demonstrated. For example, Wiener et al. (2011) leveraged the TCT to dissociate cognitive mechanisms of human path integration. Harris & Wolbers (2012) identified significantly impaired path integration performance in elderly compared to younger human participants, using a virtual reality version of the TCT. The TCT also carries promise for advancing clinical diagnoses, with TCT ability in individuals with prodromal dementia differs significantly from health age matched controls (Howett et al., 2019; Newton et al., 2024).

The utility and clinical relevance of the TCT underscores the importance of having a homologous task in a rodent model. A rodent TCT would enable maximally relevant study of the neurobiological mechanisms of path integration, of path integration ability in transgenic models, and of preclinical assessment of therapeutic interventions. Yet a rodent homologue of the TCT does not currently exist. Rodent path integration is frequently assessed using a variety of tasks including classic examples such as the Morris Water Maze and Barnes Maze (Barnes, 1979; Morris et al., 1982) alongside more recently developed tasks (Bower et al., 2005; Guerrero et al., 2023; Najafian Jazi et al., 2023). While these tasks have proven utility for the study of rodent path integration; they differ from the TCT in important ways. There remains, therefore, a requirement for a homologous rodent version of the TCT which retains the essential elements of the human path integration task of which there are several: The first element is the guided portion of the task where the animal is guided through the first two vertices of a triangular path. The unguided return portion of the task, where the animal is rewarded for returning to the first location in the absence of environmental cues (e.g. in darkness). Multiple triangle completions are performed within sessions with the goal location and triangle geometry randomly varying between trials.

Alzheimer’s disease is a particularly relevant preclinical area of research and a rodent homologue of the TCT task would advance ongoing research. One of the first areas of the brain to suffer neurodegeneration is the entorhinal cortex (Jack & Holtzman, 2013). The medial part of the entorhinal cortex has been shown to be instrumental in the ability of rats to perform accurate path integration (Gil et al., 2018; Jacob et al., 2017; Parron & Save, 2004; Tennant et al., 2018; Van Cauter et al., 2013). Spatial memory is one of the first cognitive functions to deteriorate in AD (Newton et al., 2024; Ritchie et al., 2018). Therefore, the development of sensitive and specific tests of spatial memory is potentially pivotal in the early diagnosis of the disease. Work in this area has already begun. Multiple studies have demonstrated that midlife deficits in the path integration performance of human participants, as measured by TCT, were predictive of AD risk (Howett et al., 2019; Mokrisova et al., 2016; Newton et al., 2024). Investigations into cognitive deterioration and its neural bases in early AD, are necessarily split between human and rodent studies, with the promise that insights obtained in rodents will translate to human care. Yet, translational successes have been embarrassingly rare. The high failure rate is likely a product of the low external validity of rodent models (Pound & Ritskes-Hoitinga, 2018). This once again, highlights the importance of unifying the assessment tools used across species.

To address the disparity in human and rodent path integration assessment, we describe here a novel rodent version of the TCT used in humans, that facilitates the comparison of path integration performance between humans and rodents. We demonstrate that rats were able to perform the task, reliably completing multiple triangles per session. Furthermore, using a rat model of familial AD - TgF344-AD (Cohen et al., 2013) - we evaluated the novel task’s potential for studying AD-associated spatial memory deficits.

## Methods

The final goal of the present work was to develop a rodent homologue of the human triangle completion task (TCT). For total transparency we describe the process through which a cohort of rats were trained to perform the TCT and the improvements that were made to the task setup and protocol during their training and testing. First however, we describe a finalized version of the task which incorporates these improvements.

### Rodent TCT – Finalized

Rats completed a daily 15 minute session in which they completed as many trials as possible. During a single TCT trial, rats are first guided between a series of three, wall mounted, liquid reward dispensers using LED cues. Then, cued by an audio tone, rats path integrate back to the first dispenser, in the absence of visual cues, to complete the characteristic triangular path of the TCT. This task structure mirrors that of the TCT regularly used to test human path integration.

### Apparatus

Behavioural testing was performed in a circular (55cm diameter; 50cm walls) arena surrounded by blackout curtains to occlude extra-maze visual cues (Fig. 1C). The diameter of the arena was chosen so that individual triangle leg lengths are approximately two rat body lengths, with the aim of aligning to leg-lengths used in contemporary human TCT studies (e.g. Howett et al., 2019; Mokrisova et al., 2016; Stangl et al., 2020). Twelve dispensers were mounted to the walls of the arena at regular 30⁰ intervals (Fig. 1C). Though only three are needed to form the triangle on each trial, the remaining act as lures, probing for errors. We used twelve to balance the sensitivity of the task and the technical demand of constructing and controlling the apparatus. Individual dispensers consisted of plastic rectangular housing with an infra-red LED mounted to the top pointing at the behavioural tracking camera mounted above the arena, a green LED facing the centre of the enclosure and a metal reward delivery funnel mounted to the front (see Fig. 1D). The top-mounted infra-red LED was used to indicate the dispenser status to the behavioural tracking system. The green LED was selected to be bright enough to be reliably visible to the rat but not bright enough to illuminate the enclosure. The reward delivery funnel was selected to be aluminium to discourage rats from chewing the funnel between trials. The dispenser was hung such that the funnel top was about 12 cm from the floor, requiring rats to rear to check the funnel content and thereby allow us to distinguish between when a rat generates an overt response and when it passes by the dispenser. A custom system was built to allow for the remote liquid reward delivery to each of the dispensers. A computer-controlled piston (a 100ml syringe) was used to pump reward to each dispenser individually through a series of computer controlled solenoid valves via an Arduino interface. This system was controlled by the experimenter through a custom software interface. In addition to controlling reward delivery this software also allowed the experimenter control both the illumination of dispenser LED and visible illumination through a halogen lamp mounted above the arena. We refer to this light henceforth as the house light. Infra-red (IR; 850nm) LED strips are also suspended above the arena alongside an IR camera for monitoring and recording behaviour irrespective of visible illumination. Additionally, the computer interface controlled a speaker, used to play a noisy audio tone (generated as white noise that was bandpass filtered to 2.7-3.3 kHz) for a prescribed duration. This custom reward delivery system and computer interface were built so that the experimenter and the animals need not interact during behavioural testing, reducing experimenter related behavioural confounds.

**Figure 1.**
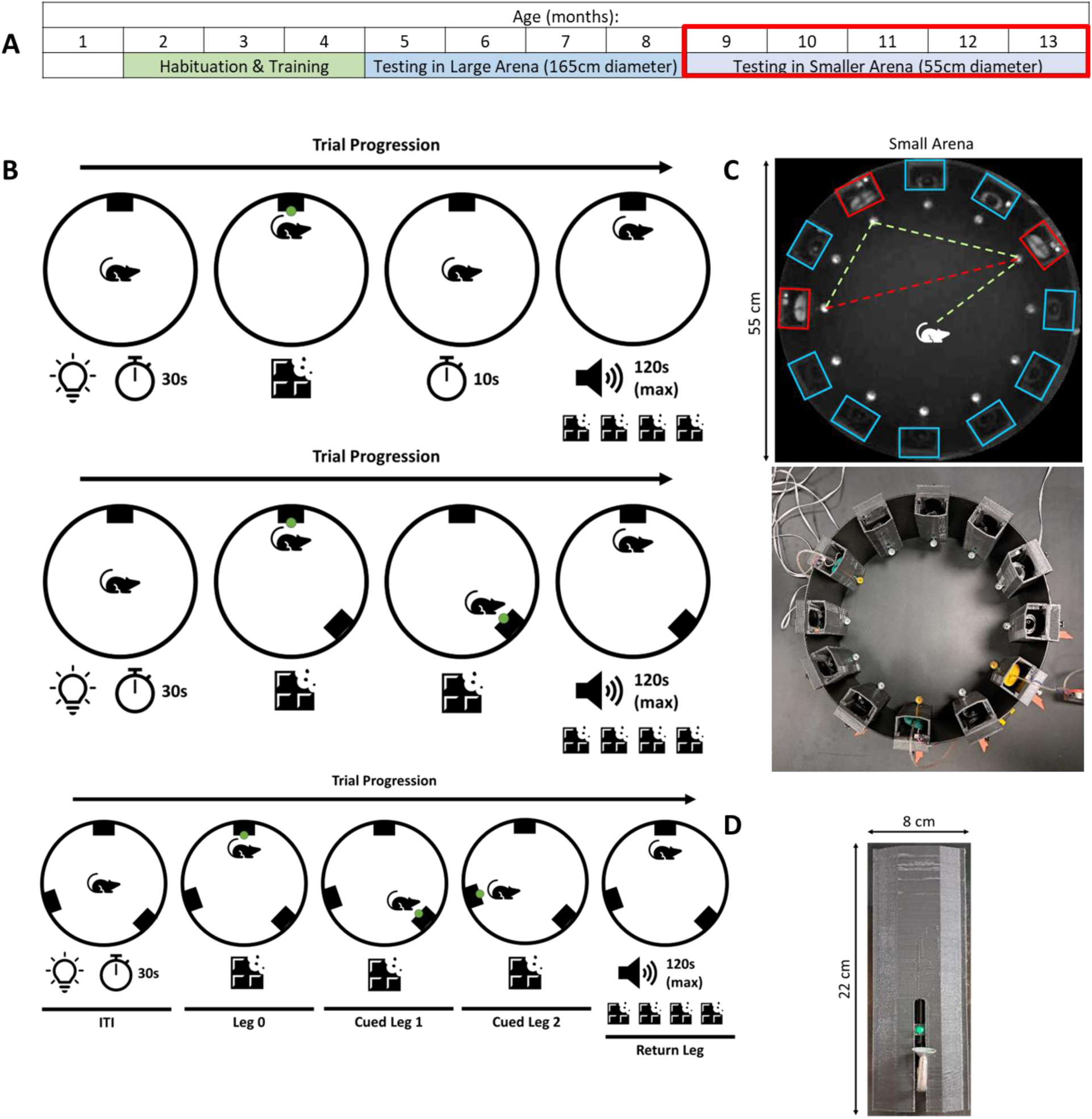
Triangle Completion Task Apparatus and Progression: (**A**) Timeline of the shaping and testing schedule that the animals underwent for the TCT. The data described here were gathered during the testing period highlighted by the red box. (**B**) Diagram representation of the three stages of shaping (rows 1-3) used to steadily introduce the mechanics of the TCT to the animals. The different epochs of the task are labelled on the final row. These include the inter-trial interval (ITI), all cued legs and the final return leg of a TCT trial. (**C**) The upper panel is a diagram demonstrating the full task where the rat runs between the three active dispensers (highlighted red) for the guided portion of the task (green dashed lines) before returning to the first dispenser visited (red dashed line). Decoy, non-target dispensers are highlighted in blue. The lower panel is a photograph of the experimental set up for the smaller arena in which the testing took place. (**D**) A photograph of a liquid reward dispenser, showing the metal dispenser funnel at the front and the green LED above the funnel.

#### Procedure

In complete darkness, rats are visually cued to approach and retrieve liquid reward from three dispensers in randomly chosen locations (of a possible 12) before an audio cue signals the availability of an increased reward at the first of the three dispensers. Consecutive illumination of the green LEDs on the front of each dispenser visually cue the rats to approach each dispenser in sequence. Once the rat retrieves liquid reward from the dispenser the LED on the current dispenser was extinguished and the LED on the next dispenser in the sequence was illuminated. When the rat completed the guided portion of the task, retrieving liquid reward from each of the three dispensers, the return phase of the trial began as a noisy audio tone was played through a speaker near the data acquisition PC to indicate to the animal the presence of an increased (4x) reward at the dispenser visited first. Note that during this return phase, the location of the dispenser is no longer indicated by the LED. The tone was played for a maximum of 2 min, during which the reward was available; after this, however, the trial was simply abandoned without reward. To promote rapid responses, the time for which the audio tone played was adjusted based on performance in the previous trial. If the previous trial was successfully completed within the time limit, the time limit for the current trial was reduced by 10s. If the previous trail was unsuccessful the current trial time limit was increased by 10 s. A lower limit was set to 15s and an upper limit was set at 2 min. Trials were separated by a 30s inter-trial interval during which the house lights were turned on. The following trial then took place using a different, randomly selected, set of three dispensers.

#### Behavioural Shaping

Due to the relatively complex nature of the task mechanics, animal behaviour was shaped in several phases prior to testing. This allowed the animals to learn the various task requirements and the sequence of events for a given trial. Across all phases, individual trials were run in a complete lack of visible light. As in testing, shaping sessions lasted 15 minutes, during which, animals performed as many trials as possible. Upon reaching a criterion of 3 trials on 2 consecutive shaping days, animals progressed to the next phase.

#### Habituation

Rats were handled for 10-15min daily by experimenters for 2 weeks prior to shaping. Rats then habituated to the reward dispensers in their home cages by hanging the dispenser from the wall of the cage. During this time, both the LED and audio tone were turned on in alternation and the dispenser baited to coincide with each. This served to train rats to associate the visual LED cue and the auditory tone cue with the availability of reward at the dispenser. Rats also habituated separately to the testing arena first in groups, along with their home cage mates, for 20 minutes and then individually for 10 minutes. This extensive habituation was performed in an attempt to manage the known anxiogenic phenotype of the TgF344-AD animals (Pentkowski et al., 2018, 2022).:

Stage 1:

In the first stage of shaping 3 dispensers were present on the walls of the arena and the animal guided to and retrieved reward from each dispenser in sequence before the audio tone was played to signal the return phase where the rat attempted to return to the first dispenser for an increased reward (4x; Fig. 1B, third row).

Stage 2:

Stage 2 consisted of the exact same procedure as stage 3 but the positions of the dispensers were randomized at the start of each session. Dispenser positions were sampled from a total of 12 possible dispenser positions distributed uniformly at regular 30⁰ intervals around the arena.

Full task:

Finally, the full task was identical to stage 4, however all dispenser positions were occupied dispensers (as seen in Fig. 1) and the three dispensers forming the triangle were pseudo-randomised between trials. The 9 dispensers not active during a given trial served as decoys.

### The Present Study – TCT Development

Here we describe the version of the TCT used in the present study, which underwent development as the study progressed.

#### Subjects

All animal procedures were conducted in strict accordance with National Institutes of Health the Indiana University Institutional Animal Care and Use Committee guidelines.

Eighteen rats were the subjects of this study. 11(9F & 2M) of these animals were F344Tg-AD, carrying mutant human amyloid precursor protein (APPsw) and presenilin 1 genes (PS1ΔE9; Cohen, 2013). 7 (1F & 6M) were age-matched wildtype (WT) littermates. TgF344-AD rats, developed by Cohen et al. (2013), exhibit AD-associated, age-dependent increases in amyloid beta and tau toxicity, along with various cognitive deficits.

Experimenters were blind to the genotype of each animal throughout behavioural testing and data scoring. Rats were bred in house. Original transgenic breeders were obtained from the Rat Resource & Research Centre (Columbia, MO) and wildtype breeders were purchased from Inotiv (Indianapolis, IN). Genotype was verified by Transnetyx using ear punches collected at postnatal day 21 during weaning.

Rats were either group or pair housed at all times and maintained on a 12h light/dark cycle in a temperature and humidity-controlled room with ad libitum access to water, and food restricted to maintain ∼90% (85–95%) of free feeding body weight. Rats were a bred in house. All work with the animals was performed during the light cycle. Daily handling began at P30.

#### Apparatus

Training and testing took place as described above with the following differences: We experimented with two other sizes of enclosure. The rats were run in a large (165cm diameter) for ∼4 months (mos. 5-8) and a small (45cm diameter) enclosure for one week before we settled on the 55cm diameter enclosure.

#### Procedure

Shaping of the animals to perform the task was done as previously described with the following minor differences:

#### Habituation

Rats were handled for 10-15min daily by experimenters for 7 weeks prior to shaping (1.5-3mo). Habituation was extended while the experimental setup was refined. Rats were not trained to associate the audio tone and the liquid reward in the home cage during habituation to the dispenser.

#### Shaping

Shaping in this initial version of the task was extended with two additional stages of shaping prior to the first stage described above in the finalised version. These stages were initially used to introduce the task mechanics in their simplest form; however were later assessed to be adding unnecessary complexity for little return in performance.

Stage 1:

A trial in the first additional stage of shaping consisted of the rat being guided to a single dispenser using the green LED (only a single dispenser present in the arena). Once the rat retrieved the reward, the LED was turned off. After a 10s pause, the unguided, return phase of the trial began as signalled by the audio tone. The audio cue was played for a maximum of 2 min, during which the reward was available, note that the audio tone length was not dynamically adjusted during this phase.

Stage 2:

During the second additional stage of shaping, two dispensers were present within the arena and rats were trained to visit each in the guided phase before returning to the first one in the return phase (Fig. 1B, second row). Note that once again the audio tone length was not dynamically adjusted at this stage.

Shaping then continued as described above in the finalized task description.

Full task:

The full task proceeded as above; however, in this initial experiment the dispenser positions remained constant within a session meaning that the rats repeated the same triangle completion within a session. The triangles were however pseudorandomised between sessions. The ability to dynamically adjust the active dispensers between trials was not yet functional in our custom reward delivery system.

### Behavioural Analysis

Animal behaviour was recorded on an infra-red camera suspended above the testing arena. Offline analyses, using DeepLabCut (DLC), tracked the position of the nose, midback and tail base (rump). Coordinates were imported into MATLAB for parsing and analysis.

#### Markerless pose estimation

Rat tracking was performed offline with DLC. A new model was built using 20 frames from each of the 18 videos from this study (360 total images) cropped to a tight circle surrounding the testing enclosure. The model was trained for about 400k iterations before it was used to analyse the training videos and 20 ‘outlier frames’ from each video were extracted based on the “jump” approach. After correcting the labels on the extracted outlier frames, the model was retrained with the 720 total frames for an additional ∼1000k iterations. DLC outputs were post-processed in MATLAB using custom scripts to drop low confidence estimates and jumps. Missing data was replaced with linear interpolation.

#### Video Annotation

Within each video, instances of specific features and events in task progression were annotated using the video recording and the DLC tracking data:

*Funnel localization:* The experimenter manually specified the position of the three active dispenser funnels using the Matlab function ginput.m. The remaining funnels were identified as peaks in luminance along the circumference of a circle drawn through these active dispenser locations. Each of the dispensers was then assigned a unique numerical label depending on its location in the arena.

*LED localization:* The experimenter also specified regions of interest around each of the infra-red LEDs atop the active dispensers (IR-ROIs).

*Dispenser LED and house light status:* The video was processed frame-by-frame to extract the time-varying mean luminance within the IR-ROIs and across the whole video frame. Fluctuations in the whole frame luminance were thresholded to track when the overhead house light was off versus on, marking the inter-trial intervals. Fluctuations in the IR-ROIs were also thresholded to mark epochs when the LEDs were illuminated.

*Funnel ‘pass-bys:’* The behaviour was parsed to identify the time of arrival and departure from each funnel. This was done by comparing the distance between the rat’s nose and each of the dispensers for each frame of the video. A dispenser pass-by occurred when the rat’s nose passed within 25 pixels (5.5cm) of a dispenser funnel and remained there for at least 0.5 seconds. Pass-by events separated by less than a second were considered the same event. A pass-by could have included a nose-poke or not.

*Funnel nose-pokes:* A nose-poke is when a rat engages with a dispenser enough to occlude some of the funnel from the view of the overhead camera. Because this requires the rat to rear, it is used as an indicator of overt responding. Each frame within a cropped video of each dispenser funnel was compared to an image of the same funnel captured in a reference frame using MATLAB function immse.m. When the rats nose entered the funnel the luminance of the pixels within the frame changed relative to the reference frame allowing for the nose-poke to be detected.

#### Behavioural Epoch Definition

Behavioural epoching was done to mark the start and stop times of the key phase of each trial: Leg 0, as the rat approaches the starting point of the triangle (i.e., dispenser 1); Cued Leg 1, as the rat moves from dispenser 1 to dispenser 2; Cued Leg 2, as the rat moves from dispenser 2 to dispenser 3; and Uncued Return Leg, as the rat returns from dispenser 3 back to dispenser 1 (see Fig. 1B final row). The start of Leg 0 was defined by the time that the LED on the first dispenser turned on. Leg 0 ended when the rat nose-poked the first dispenser. The beginning of all remaining epochs were defined by the end of the pass-by window of the previous dispenser, indicating that the rat left the immediate proximity of that dispenser. The end of each epoch was defined by a nose-poke at the target dispenser. In the case of the return epoch – if the rat did not return during the audio-tone-indicated time limit the trial end was defined by the house light turning on (at the beginning of the inter-trial-interval) and the trial marked as incomplete.

#### Quantification of TCT Performance

TCT performance was quantified using path length, latency, thigmotaxic index, tortuosity index and incorrect dispenser visits.

*Path length* quantifies the distance travelled in a leg. Path length was calculated using the rump coordinate from DLC tracking, instead of the default nose coordinate, to prevent non-locomotor head movements (e.g., grooming) from artificially extending the path length.

*Latency* is the duration of each leg, irrespective of how the time was spent.

*Thigmotaxic index* scored the tendency of the animal to remain against a wall during the leg. This was scored as:

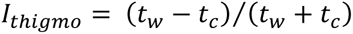

Where *t_c_* and *t_w_* reflect the time spent in the enclosure center and along a wall, respectively, such that negative values reflect more time spent near the walls versus in the enclosure center. The rat was scored to be along a wall if it was within ∼4cm of the wall. Note, in practice, this was achieved with a threshold of 12cm from the apparent position of the upper rim of the enclosure as viewed from the top-down camera.

*Tortuosity index* scored the ratio of the path length taken to the shortest possible path length. A value of one indicates the shortest possible path and larger values reflect how much longer the path length was than the minimum distance (e.g., 2 means twice the shortest path length).

*Incorrect dispenser visits* were defined by visits to dispensers other than the target during a given epoch of the TCT trial.

### Statistical Analysis

One-sample t-tests were used to assess if the proportions of epochs where correct and erroneous dispenser locations were visited was above chance levels. Only epochs that were part of completed trials were used for this analysis as rats were given a maximum of 15 minutes to complete as many trials as possible, there were occasions on which a trial was terminated before it was finished. Mann-Whitney U tests were used to assess differences in TCT performance between the transgenic and wild-type animals, as measured by latency, path length, tortuosity index and thigmotaxic index. Differences in head-scan index were assessed using an independent samples t-test.

## Results

### WT and Tg-F344AD rats are able to learn the task mechanics

All animals were trained in the TCT initially in a large (168cm) arena before testing commenced in a smaller arena (Fig. 1A; 55 cm). The diameter of the arena was reduced to better match the relative scale at which humans usually perform the TCT. Tracking data was generated from videos, recorded from an infra-red camera suspended above the arena, using the behaviour tracking software DeepLabCut. This allowed us to plot, parse and analyse the animal’s trajectories in Matlab (Fig 2A). Animals completed a total of 64 days of testing in the smaller arena (Fig. 1A) resulting in 1100 videos. Automated trajectory analyses were used to parse the behaviour in each video – trajectories in 840 videos were successfully parsed and analysed this way.

**Figure 2.**
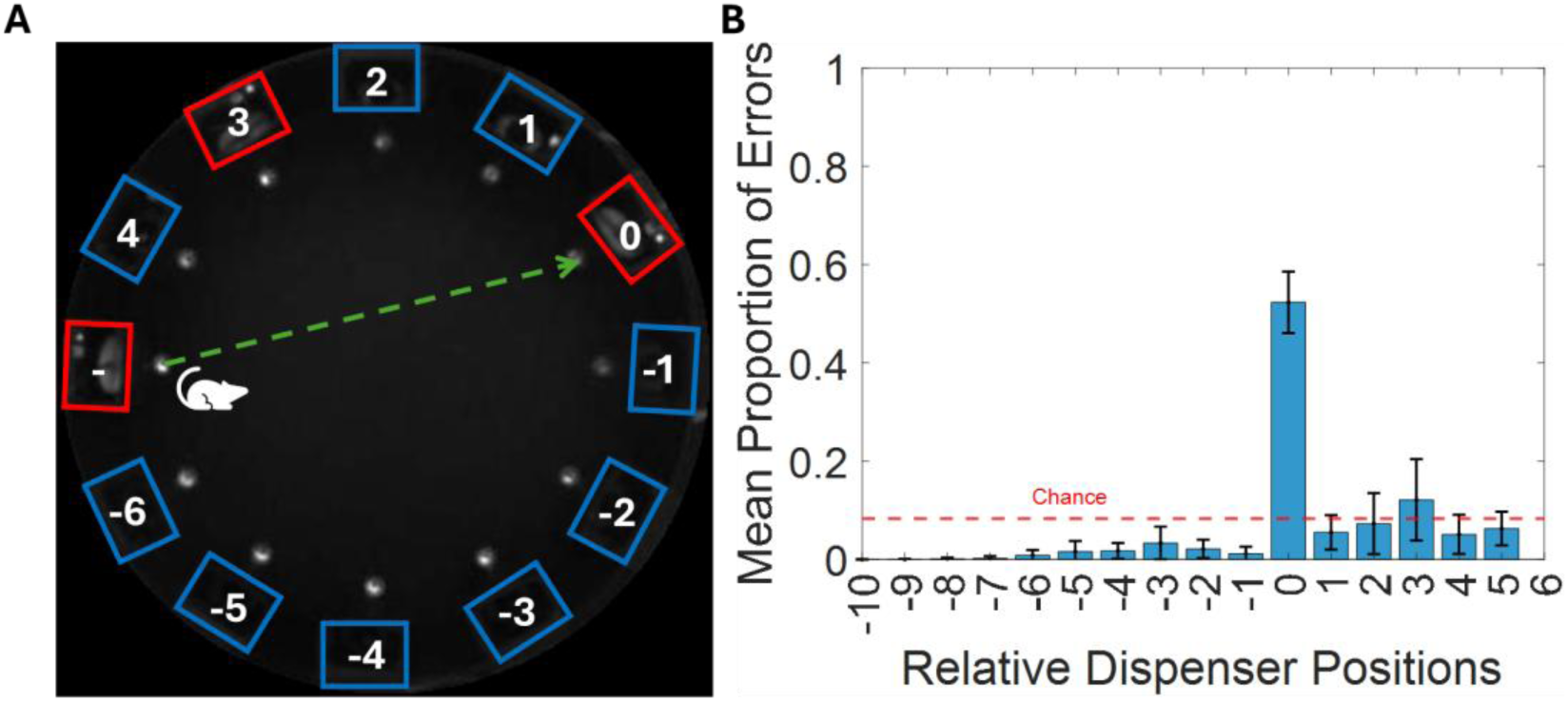
Rats Comprehend Triangle Completion Task Mechanics: (**A**) Diagram demonstrating the method by which dispensers were labelled in order to assess the distribution of visits during each epoch. The target dispenser is labelled as 0 and non-target dispensers are labelled by their position relative to the target. Dispensers on the shorter arc between the start and the target were labelled with positive integers and those on the longer were labelled with negative integers. (**B**) Bar graph showing the proportion of visits made to each of the dispenser positions, described in A, during the guided epochs of the task. The red dashed line denotes the chance level (0.083).

The task was split into four distinct epochs. The first three epochs formed the guided portion of the task, where animals approached 3 dispensers in sequence, cued by a green LED, to retrieve liquid reward. We first examined the performance of both WT and TgF344-AD animals in these guided epochs to allow us to assess their comprehension of the task mechanics. For each epoch we measured the proportion of occasions on which rats visited the target vs non-target dispensers. To assess the distribution of these visits we labelled each of the non-target dispensers using their position relative to the target dispenser (labelled as 0) and the starting location of that epoch (Fig 2A). Non-target dispensers positioned on the shorter arc between the start and target dispensers were labelled with positive integers; while those positioned on the longer arc were labelled with negative integers. Of a choice of 12 possible dispensers, the animals exclusively visited the correct, target, dispenser on 53% of guided epochs (Fig. 2B). This performance is significantly greater than chance (t_(17)_=29.99, p<.001). Rats did not visit any incorrect dispenser with a frequency significantly greater than chance. This finding is consistent with the rats maintaining good comprehension of the task mechanics. Notably, however, the fact that animals visited non-target dispensers on 47% of guided epochs indicates a considerable amount of non-task directed exploration, and therefore a possible lack of motivation.

### WT and Tg-F344AD animals are able to perform path integration in the Triangle Completion Task

Following the three guided epochs, the final epoch within a trial, the return epoch, was used to assess path integration performance. This leg of the TCT required the animals to navigate back to the first dispenser location in the absence of external landmarks (i.e. in complete darkness), thus requiring them to path integrate (Fig 3A, 4^th^ column highlighted green). We first assessed their performance by measuring the proportion of return legs on which rats visited the target vs non-target dispensers. As described above, a visit to a dispenser was defined by the rat rearing and poking it’s nose into the dispenser funnel. Simply passing close to the dispenser was not counted as a visit. We found that, on average, both TgF344-AD and WT rats exclusively visited the target dispenser on ∼80% of trials (Fig. 3B) - this performance was significantly greater than chance for both groups (WT: t_(6)_=29.03, p<.001; TG: t_(10)_=32.85, p<.001). Furthermore, neither group visited any of the non-target dispensers with a frequency above chance levels, during the return epoch. This finding suggests that these animals understood the objective of the task and were motivated to visit the target dispenser exclusively. Furthermore, the relative positions of visits to non-target dispensers were evenly distributed, indicating no systematic errors in their path integration or task comprehension in either group (Fig. 3B).

**Figure 3.**
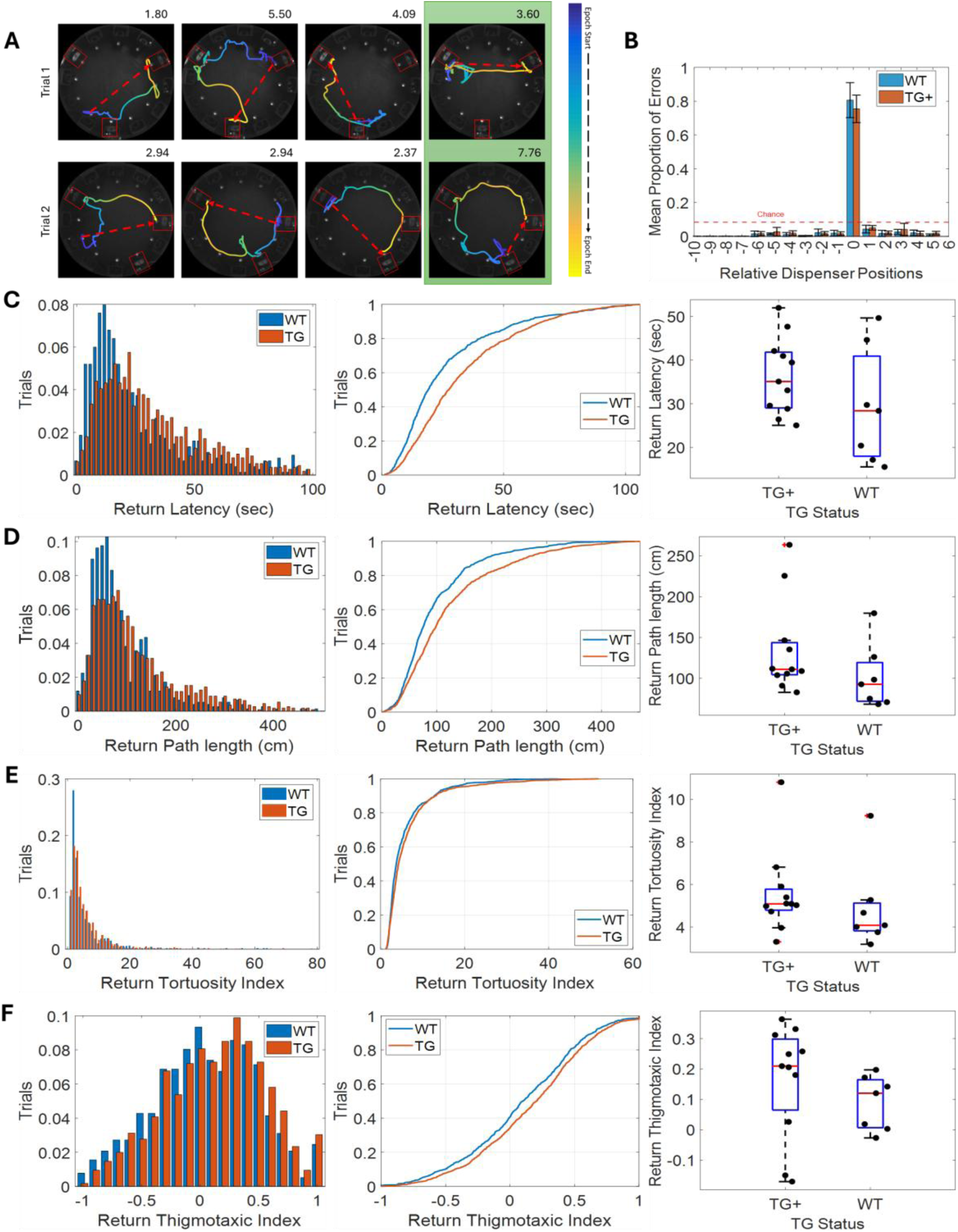
Both Wild-Type and Transgenic rats perform the Triangle Completion Task: (**A**) Example trajectories from two triangle completion task trials (top and bottom rows respectively). Columns 1-3 contain the 3 guided epochs and the 4^th^ column (highlighted in green) contains the return epochs for each trial. The red dashed arrow indicates the shortest possible path for each epoch and the value at the upper-right corner of each plot is the tortuosity index for that epoch. (**B**) Bar graph showing the mean proportion of visits made to each of the dispenser positions during the return epochs. The red dashed line denotes the chance level (0.083). (**C, D, E, F**) Graphs showing return latency, path length, tortuosity index and thigmotaxic index respectively. The left hand column contains histograms; the middle column contains cumulative density plot and the right hand column contains box and whisker plots. These graphs all compare the performance of WT and TgF344-AD animals during the return epoch of the TCT for each measure.

Next, we assessed path integration performance through measuring return epoch latency, path length and tortuosity index (Fig 3C, D, E). These measures demonstrated no significant differences in the median performance of the two groups, demonstrating that WT and TgF344-AD animals had equivalent proficiency in this task (TG+ vs WT: Latency - U_(16)_=25, *p=.238*; Path Length – U_(16)_=20, *p=.103*; Tortuosity Index - U_(16)_=24, *p=.204*; Thigmotaxic Index - U_(16)_=19, *p=.085*). There was, however, a consistent trend towards the performance of TgF344-AD animals being marginally impaired compared to their WT counterparts, especially in the case of latency and path length (Fig 3C, D). Examining the trajectories from both WT and TgF344-AD animals suggested their behaviour was thigmotaxic (i.e. running next to the walls of the arena; Fig 3F). Both WT and TgF344-AD animals spent much of their time next to the walls of the arena, indicating a significant level of anxiety driven behaviour. Furthermore, there was, once again, a trend towards the TgF344-AD animals having a greater thigmotaxic index compared to their WT littermates, suggesting their behaviour was more anxiety driven.

The dispensers forming the triangle changed between sessions; within a single session the rats performed multiple trials completing the same triangle. It could, therefore, be argued that following the first trial, the requirement for path integration in this task was lost for the remainder of the session. To account for this, we next examined path integration performance using only the first trial from each session. In line with previous analyses, the performance was very similar between WT and TgF344-AD animals (as measured by latency, path length, tortuosity index and thigmotaxic index) with a trend towards marginally impaired performance in the TgF344-AD animals once again across most measures (Fig 4; TG+ vs WT: Latency - U_(16)_=22, *p=.147*; Tortuosity Index - U_(16)_=21, *p=.124*; Thigmotaxic Index - U_(16)_=27, *p=.317*) however the increase in path length was statistically significant (U_(16)_=15, *p=.038; Fig 4B*).

**Figure 4.**
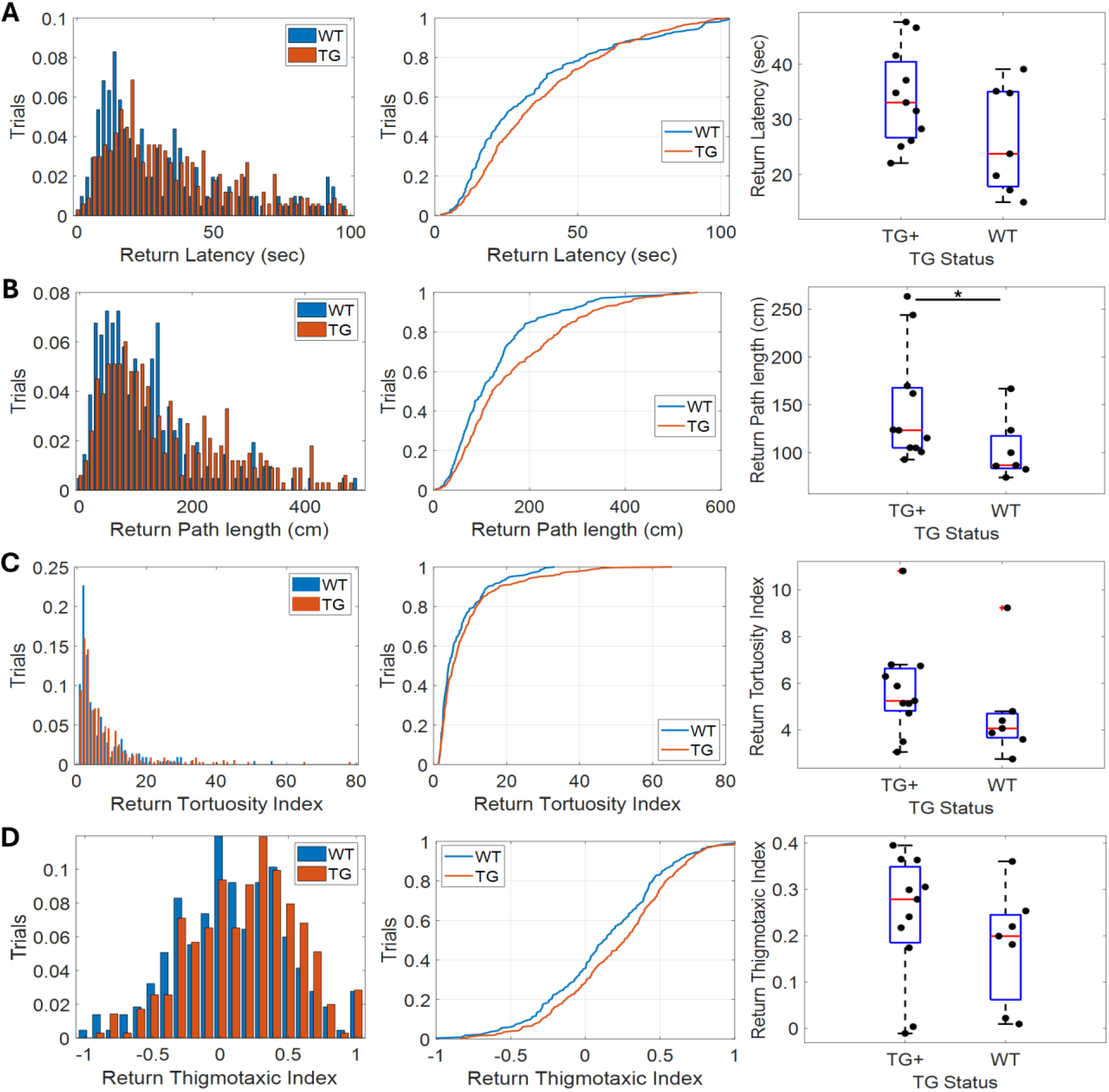
TCT performance is unchanged if only using data from 1^st^ trial of each session: (**A, B, C, D**) Graphs showing return latency, path length, tortuosity index and thigmotaxic index respectively. The left hand column contains histograms; the middle column contains cumulative density plots and the right hand column contains box and whisker plots. These graphs all compare the performance of WT and TgF344-AD animals during the 1^st^ return epoch of each TCT session for each measures. * indicates p<.05

While our data suggests that the path integration performance is similar between the TgF344-AD and WT animals there was, however, some indication that the two groups may not have been performing the task in the same way. We assessed the extent of lateral head-scanning in the two groups, during all trials, using a head-scan index (path length measured by nose tracking divided by path length measured by rump tracking). Wildtype animals exhibited significantly greater lateral head-scanning than their TgF344-AD counterparts (Fig 5, t_(16)_=4.15, p<.001).

**Figure 5.**
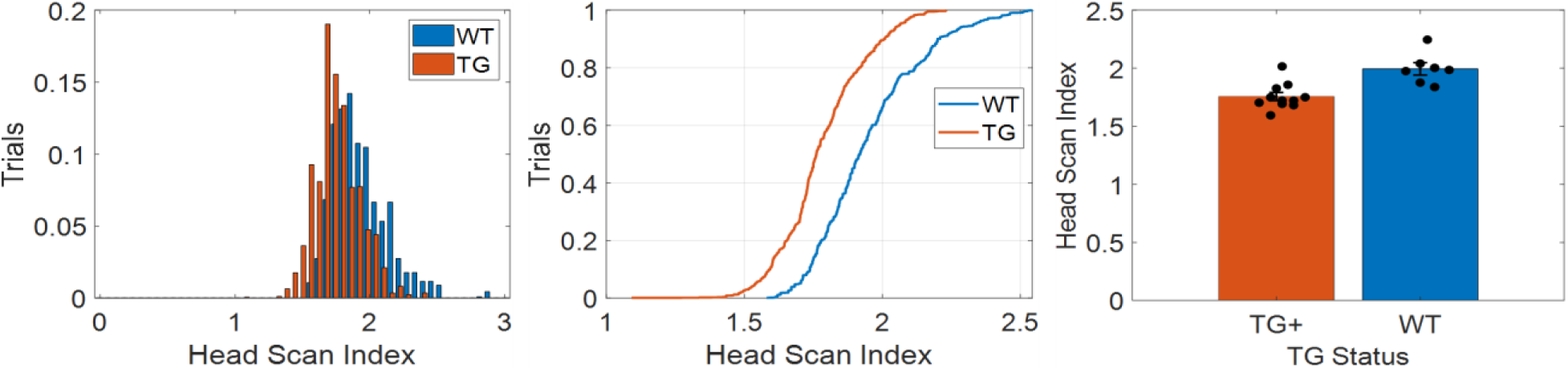
WT animals exhibit significantly more lateral head-scanning than TgF344-AD animals: The panels from left to right contain a histogram, cumulative density function and a bar graph of head-scan index, respectively.

## Discussion

Path integration has been studied extensively across both humans and rodents; however, the tasks used to assess it differ significantly between species. Homologous tests of path integration in humans and rodents are important as rodents are routinely used as preclinical models. In the present study we developed a novel rodent version of the TCT which retained the essential elements of the human variant of the task. Rats demonstrated that they were able to comprehend and perform the task, whilst the comparisons between TgF344-AD animals and their WT littermates revealed that both groups performed the task with similar proficiency. There was a consistent trend towards an impairment in the TgF344-AD animals’ performance. Additionally, we observed evidence of a significant difference in the amount of lateral head-scanning between the genotypes, suggesting that they used different strategies. Together, these results show that this rodent variant of the TCT, is a functional homologue to the human TCT.

Rodent path integration is frequently assessed using a variety of tasks with proven utility. Classic examples of these tasks are the Morris Water Maze and Barnes Maze (Barnes, 1979; Morris et al., 1982). The Morris Water Maze involves placing the animal into a circular pool filled with opaque water. The animal must then use path integration to find its way to a small platform, submerged just under the surface, in order to escape the water. Similarly, the Barnes maze involves placing the animal on a circular platform featuring a number of evenly spaced holes around the perimeter. The animal must find its way to the hole which leads to an escape box. Both of these tasks have proven utility in the assessment of path integration in rodents. Their use of escape behaviours; however, makes performance comparisons between these tasks and the TCT challenging. Furthermore, the use of escape as motivation in these tasks may result in stress driven behaviour, and the disruption of optimal spatial memory performance (Sandi & Pinelo-Nava, 2007; Schwabe & Wolf, 2010).

In contrast to classic path integration tasks, more recently developed spatial memory tasks bare a closer resemblance to the TCT (Bower et al., 2005; Guerrero et al., 2023; Najafian Jazi et al., 2023). In one task, dubbed AutoPI, a mouse searches a circular arena for a randomly placed lever which they operate in order to receive a reward upon returning to a home box (Najafian Jazi et al., 2023). Advanced apparatus and custom software are utilized to automatically progress each trial allowing for many trials per session.

This impressive task is a useful test of path integration in rodents. The use of a home box, however, introduces an element of escape behaviour as rodents prefer enclosed spaces to the open field of the testing arena. This differentiates this task from the TCT which uses positive motivation instead. Another spatial memory task which bares resemblance to the TCT was originally described by Bower et al. (2005) and more recently refined by Guerrero et al. (2023). Rats remain in an open field throughout testing and are guided between different locations around the perimeter of a circular platform using LED cues. Rats then return to one of two constant reward locations, depending on the sequence of locations visited during the trial, to receive a reward. This task has primarily been used to study sequential memory in hippocampus and has yet to be used to study path integration. Thus, the task setup and procedure require some modification in order to assess path integration in a manner similar to the TCT. For example, pseudo-randomly changing reward locations in place of the two constant reward locations in the present version of the task. To effectively study path integration, the overlapping paths and fixed reward locations in the current task design may inadvertently introduce interference from olfactory cues. It is important to note that the differences between these tasks and the TCT do not call into question their validity or utility. These differences simply illustrate the remaining requirement for a TCT homologue in rodents.

The lack of clear path integration performance differences between 9-13 month old TgF344-AD rats and their WT littermates in this study contrasts with prior work. For example, TgF344-AD rats show significantly impaired Morris Water Maze performance as early as 6 months of age (Bac et al., 2023; Berkowitz et al., 2018; Bernaud et al., 2022) and exhibit spatial memory deficits in an active place avoidance task (Chaudry et al., 2022). As already noted above, behaviour in these tasks, however, is motivated by escape. Thus, it is possible that previously observed impairments in TgF344-AD rat performance could have been exaggerated by transgene related differences in escape motivation. The use of non-appetitive motivation differs substantially from the positive reinforcers used in human path integration assessments and from what we used in our variant of the TCT. While TgF344-AD rats have also been shown to have impaired performance in an appetitively motivated T-maze based spatial working memory task (Saré et al., 2020) this task did not assess path integration ability. Thus, the behavioural deficits observed to date in TgF344-AD rats may not be strongly related to human AD patient path integration ability. This further underscores the importance of establishing a rodent homologue for the human path integration assessment tools as we have done here.

Path integration deficits found in human patients using the TCT (Howett et al., 2019; Mokrisova et al., 2016; Newton et al., 2024) were not perfectly replicated in the present study. Path integration deficits have been observed in humans exhibiting mild cognitive impairment (MCI) as well as those with the APOE4 allele. The path integration deficits we observed in TgF344-AD animals in the present study, however, were minimal. This disparity may be explained by several differences between the present study and those studying preclinical AD patients. The humans assessed by Newton et al. (2024) were APOE4 allele carriers, the allele associated with the greatest risk of developing sporadic AD, unlike the TgF344-AD rats used in the present study. Additionally, these APOE4 carrying humans exhibited a average angular error of less than 10 degrees. Given that the dispensers used in the present study were placed at 30 degree spacing, it is possible that the apparatus in the present study did not have the angular resolution to detect this difference in angular error. This may be remedied in future by the addition of more dispensers. Howett et al. (2019) found that amnesic MCI patients had significantly greater distance error compared to controls. The task in the present study however, is not sensitive to exclusive distance errors as all dispensers were mounted to the perimeter of the arena. There may also be remedied in future by the addition of dispensers to locations at different distances from the arena perimeter.

The similar performance of the TgF344-AD and WT rats, however, enables better controlled comparison of the genotypes with respect to other dimensions. For example, in a simple analysis of their behavioural structure, we observed that the TgF344-AD exhibited significantly less lateral head scanning than the WT rats. The difference in lateral head scanning suggests different exploratory strategies between animals and, thus, potentially different hippocampal encoding dynamics (Layfield et al., 2023; Monaco et al., 2014). The lack of gross differences between the groups leave it possible to make compare neurophysiological differences between the genotypes in future work. For example, grid cells are hypothesized to support path integration (Gil et al., 2018) but lose tuning in rodent AD models (Jun et al., 2020; Kunz et al., 2015; Newman et al., 2014; Ying et al., 2022).

One limitation of the current study was the relatively few trials completed per 15 minute testing session. We attribute this to limited motivation in the current animals. We utilized food restriction in an attempt to incentivise good performance, however, we found these animals, both TgF344-AD and WT littermates, challenging to motivate. Our animals would become under-conditioned before any improvement in motivation was observed. Tournier et al. (2021) observed similar abnormalities in these animals, finding decreased motivation and lacking response to liquid reward. Liquid reward was the mechanism for reward delivery in the present study. A further possible explanation for low trial completion is the observed increase in anxiety levels in these animals, which pre-dates cognitive decline (Pentkowski et al., 2018, 2022). While we took steps to reduce anxiety in our animals, it is possible that we were unsuccessful in entirely mitigating this factor. The fact remains, however, that despite the idiosyncrasies of this particular AD model, these rats were able to comprehend and perform the TCT. It follows, therefore, that with more easily motivated animals, the capacity of this task to detect specific differences in path integration performance may be increased. This highlights the requirement for an AD rat model better suited to cognitive behavioural testing, without the motivational and anxiety related features of the TgF344-AD model.

In conclusion, the present study demonstrated that rats are able to perform a homologous path integration task to that performed in humans; facilitating the future comparison of human and rodent path integration performance. This enhanced intra-species comparability makes this rodent TCT suitable for the reverse-translational study of the deteriorating neurophysiology underlying path integration deficits in AD.

## Notes

**Conflict of Interest Disclosure:** No conflicts of interest to disclose.

**Funding Statement:** This work has been supported by the National Institutes of Health (RO1AG076198 to EN) and Hutton Honours College research grants (to S Rehman, IC, OK, L Goldman) and Indiana University office of research integrated freshmen learning experience award (S Rehman, S Ramlo). This research was supported in part by Lilly Endowment, Inc., through its support for the Indiana University Pervasive Technology Institute and in part by Shared University Research grants from IBM, Inc., to Indiana University.

### Competing Interest Statement

The authors have declared no competing interest.

